# Fear-conditioning to unpredictable threats reveals sex differences in rat fear-potentiated startle (FPS)

**DOI:** 10.1101/2023.03.06.531430

**Authors:** Valentina Olivera-Pasilio, Joanna Dabrowska

## Abstract

Fear-potentiated startle (FPS) has been widely used to study fear processing in humans and rodents. Human studies have shown higher startle amplitudes and exaggerated fear reactivity to unpredictable vs. predictable threats in individuals suffering from post-traumatic stress disorder (PTSD). Although human FPS studies often use both sexes, a surprisingly limited number of rodent FPS studies use females. Here we investigate the effects of signal-threat contingency, signal-threat order and threat predictability on FPS in both sexes. We use a classic fear-conditioning protocol (100% contingency of cue and shock pairings, with forward conditioning such that the cue co-terminates with the shock) and compare it to modified fear-conditioning protocols (70% contingency; backward conditioning; or cue and shock unpaired). Although there are no sex differences in the startle amplitudes when corrected for body weight, females demonstrate higher shock reactivity during fear-conditioning. Both sexes demonstrate comparable levels of cued, non-cued, and contextual fear in the classic FPS but females show reduced fear discrimination vs. males. Fear-conditioning with 70% contingency or backward order (cue co-starts with shock) induces similar levels of cued, non-cued, and contextual fear in both sexes but they differ in contextual fear extinction. Lastly, a prominent sex difference is uncovered following unpredictable fear-conditioning protocol (cue and shock un-paired), with females showing significantly higher startle overall during the FPS recall, regardless of trial type, and higher contextual fear than males. This striking sex difference in processing unpredictable threats in rodent FPS might help to understand the mechanisms underlying higher incidence of PTSD in women.

**Highlights:**

- Male and female rats have comparable startle amplitudes when corrected for body weight
- Female rats show higher foot-shock reactivity than males during fear-conditioning
- Female rats show reduced fear discrimination vs. males in the classic FPS
- Reversed signal-threat order increases contextual fear in both sexes
- Exposure to unpredictable threats increases startle in general and contextual fear only in females

## Introduction

Post-traumatic stress disorder (PTSD) is classified by the fifth edition of *Diagnostic and Statistical Manual of Mental Disorders* (5th ed.; DSM–5; American Psychiatric Association, 2013) as a Trauma-and Stressor-Related Disorder. On a behavioral level, PTSD is characterized by increased vigilance, heightened startle reactivity, hyperarousal, generalization of fearful stimuli, and failure to respond to safety cues (Breslau, 2009; Grillon et al., 2009; Jovanovic et al., 2010a). Women have increased probability to suffer from anxiety disorders and PTSD than men, even when controlling for frequency and severity of the traumatic events (Kessler, 1995; for reviews see Tolin and Foa, 2006; Olff et al., 2007).

Fear conditioning has been widely used to study fear learning and expression in animal models and human studies. It involves presentations of a discrete short-duration cue (conditioned stimulus, CS), which co-terminates with an aversive stimulus such as a foot shock or air puff (unconditioned stimulus, US). Following fear memory acquisition and consolidation, presentation of the CS alone elicits fearful behavior (fear recall and expression) (Fanselow, 1980; Maren, 2001; Grillon, 2002; Fanselow and Poulos, 2005; LeDoux, 2014). In addition, sensory representations of the environment where the conditioning takes place forms an associative memory with the US, and therefore re-exposure to the context itself manifests in contextual fear (Rudy and O’Reilly, 1999; Rudy et al., 2004; Maren et al., 2013). Freezing is the most commonly measured behavioral output of fear expression in rodents. In contrast, fear-potentiated startle (FPS) has been used to study fear processing in both animal models (Walker and Davis, 2002; Missig et al., 2010; Ayers et al., 2011; Moaddab and Dabrowska, 2017) and human studies (Bradley et al., 1993; Lindner et al., 2015; de Haan et al., 2018; Szeska et al., 2021). FPS has also been a useful tool to study fear processing in PTSD patients (Jovanovic et al., 2010a, 2010b), who consistently show greater FPS, manifested as higher startle responses overall (Morgan et al., 1995), impaired fear discrimination and inhibition (Jovanovic et al., 2010a), and disproportionately higher fear reactivity to unpredictable versus predictable threats (Grillon et al., 2009), in comparison with non-PTSD controls. In FPS, the fear conditioning principle remains identical as described above, whereas fear expression is measured as a potentiation of the acoustic startle reflex (ASR) in response to a sudden acoustic stimulus. ASR itself is scored as an amplitude of an eyeblink reflex in humans (Grillon et al., 1991) or an amplitude of a whole-body jump in rodents (Davis et al., 1993). During FPS recall, the ASR is significantly potentiated by a discrete cue (or context), that has been previously paired with the US during fear-conditioning (Walker and Davis, 2002).

Because of the rapid nature of the ASR reflex (less than 200 msec from the ASR-eliciting noise onset), the FPS provides much-needed precision and temporal resolution to dissect out various cued and non-cued components of fear recall. Therefore, in addition to cued fear responses (ASR potentiation during cue presentation), FPS can also measure an anticipatory fear or anxiety when the measured ASR is higher at specific time points when rodents or humans anticipate receiving a shock (when matching the exact time points of the original fear-conditioning) (Davis et al., 1993). Furthermore, a potentiation of the ASR observed between cue presentations during FPS recall, which does not occur until after the cue is being presented (first reported by Walker and Davis (2002)), indicates that the FPS can also measure a form of cued fear generalization. Later studies referred to this FPS phenomenon as a ‘background anxiety’ (Missig et al., 2010; Ayers et al., 2011) or non-cued fear (Moaddab and Dabrowska, 2017; Janeček and Dabrowska, 2019; Martinon et al., 2019). Studies also showed that the background anxiety/non-cued fear is independent from contextual fear (Missig et al., 2010; Ayers et al., 2011).

In classic fear conditioning, as described above, paired CS-US are presented with 100% contingency, such that every CS is reinforced by the US. However, conditioned responses to CS-US with a mixed, un-paired or partial reinforcement history renders the CS uncertain or ambiguous as it is less predictable of the expected outcome (Goode et al., 2019, 2020; Glover et al., 2020; Urien and Bauer, 2022). Although FPS in humans is commonly used with varying protocols (different CS-US contingencies, predictability or order of the US) (Nelson and Shankman, 2011; Abend et al., 2022; Jovanovic et al., 2022), according to our knowledge, there are no comparative studies to date that investigated the effects of variable US predictability or contingency in rodent FPS. Furthermore, although FPS has been used in rodents, most of the studies have focused on males with a few recent exceptions (de Jongh et al., 2005; Zhao et al., 2018; Voulo and Parsons, 2019; Carvalho et al., 2021; Russo and Parsons, 2021). Considering that PTSD and anxiety-related disorders are twofold more common in women than in men (Tolin and Foa, 2006; Breslau, 2009), and higher fear reactivity to unpredictable threats is one of the hallmarks of PTSD in humans (Grillon et al., 2009), in this study we investigated how signal-threat contingency, predictability or order during fear conditioning affects FPS expression and extinction in both sexes. Specifically, we investigated the effects of reduced CS-US contingency (70% vs. 100%), an absence of CS-US predictability (unpaired vs. paired), and reversed CS-US order (backward vs. forward) on ASR baseline, reactivity to foot-shocks during fear conditioning, expression and extinction of cued, non-cued, contextual fear, and fear discrimination during FPS, and we directly compared them between male and female rats.

We report that despite females demonstrating significantly higher reactivity to foot-shocks during fear conditioning and reduced fear discrimination in comparison to males, there are no sex differences in cued, non-cued, or contextual fear recall in the classic FPS. The most prominent sex difference is uncovered following fear-conditioning to unpredictable threats (CS-US un-paired). Here, female, but not male, rats show a persistent increase in ASR overall during repeated FPS tests and a significantly higher contextual fear. This striking sex difference in processing unpredictable threats in rodent FPS might help to uncover the mechanisms underlying the higher incidence of PTSD in women.

## Experimental procedures

### Animals

Female and male Sprague-Dawley (SD) and Wistar rats (Envigo, Chicago, IL) were housed in groups of 3-4 per cage on a 12 h light/dark cycle (light 7 am to 7 pm) with free access to water and food. Upon arrival, rats were habituated to this environment for a week before experiments began. Experiments were performed in accordance with the NIH guidelines and approved by the Animal Care and Use Committee at RFUMS. A total of 163 male and female rats were used for the experiments. The average age for all rats was 56 days old (8 weeks) with average body weight of 273.3 ± 37.7 grams for males and 201.2 ± 29.4 grams for females.

### Acoustic Startle Response (ASR) and Fear-Potentiated Startle (FPS)

ASR and FPS were tested in SD male (n=32) and female rats (n=21). In order to directly compare FPS between male and female rats, absolute startle amplitudes were corrected for 100 g of body weight. Rats were tested in Plexiglas enclosures inside sound attenuating chambers (San Diego Instruments, Inc., CA), as described before (Moaddab and Dabrowska, 2017; Martinon et al., 2019). ASR was measured during 30 trials of startle eliciting white-noise bursts (WNB, 95dB) on day 1 (chamber and startle habituation), and day 2 (baseline pre-shock test). On day 3 (fear conditioning), rats were exposed to 10 presentations of 3.7 s cue light (conditioned stimulus, CS), each co-terminating with 0.5 s foot shock (unconditioned stimulus, US; 0.5 mA) in context A. On day 4, rats were tested for recall of cued and non-cued fear in context B, where ASR was first measured alone (10 trials), followed by an additional 20 trials, during which ASR was measured in the presence of the cue (light+noise trials) or in the absence of the cue (noise only trials), presented in a pseudorandom order. Context B presented altered environmental cues than context A, such as an absence of steel grid bars used for conditioning, different disinfectant used for cleaning (ethanol 70% instead of peroxide), and a different experimenter performed the testing. On day 5, rats were tested for contextual fear recall, where ASR was measured in the original context A with no cue presentations (**Fig. 1A**).

**Figure 1.**
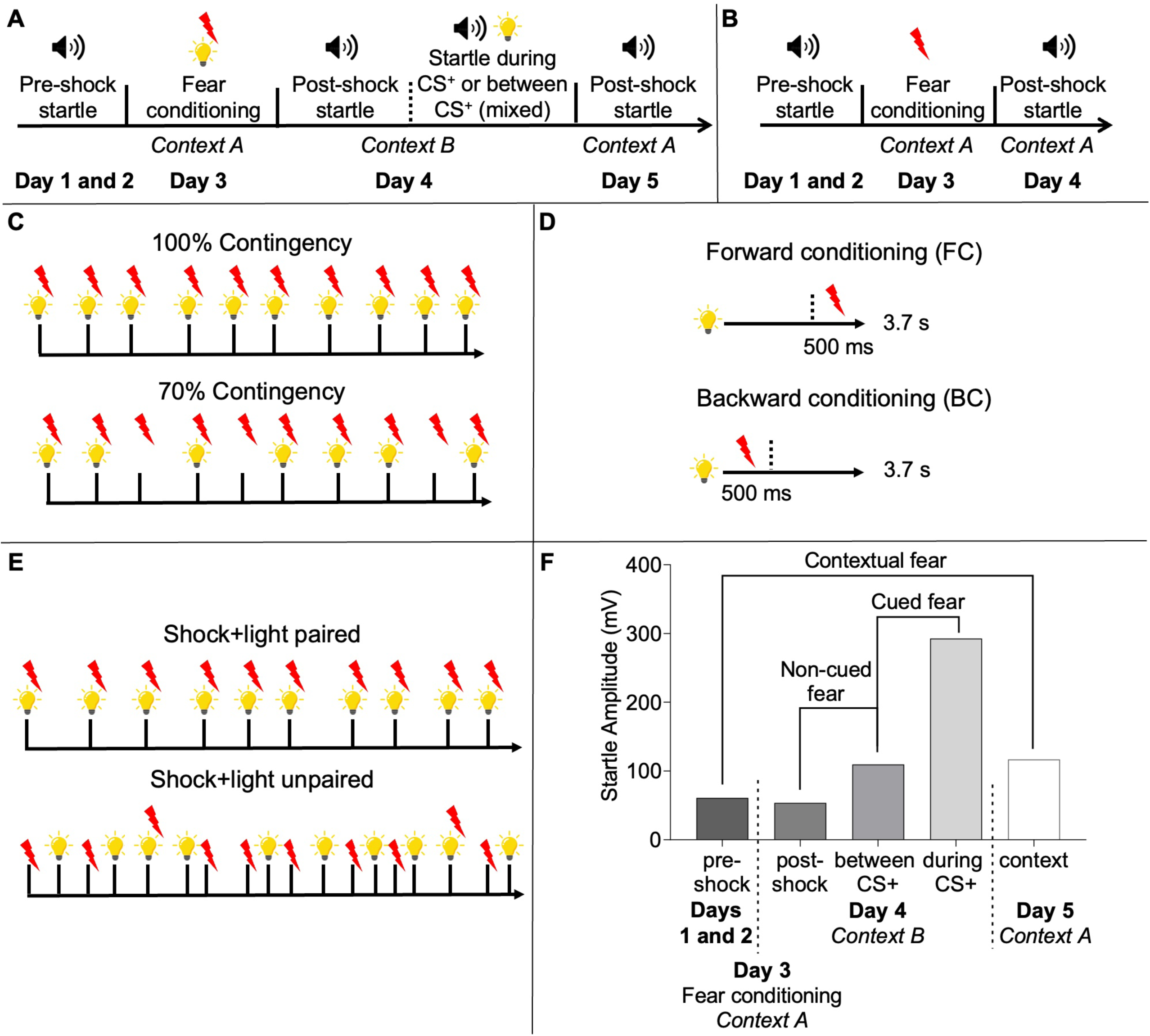
Schematic diagram of the classic and modified Fear Potentiated Startle (FPS) protocols. **A)** Classic cued fear FPS protocol timeline. All control groups are conditioned using FPS protocol with 3.7 s cue light co-terminating with 0.5 s foot-shock (100% contingency, forward conditioning, and shock+light paired). **B)** Classic contextual fear FPS protocol timeline. **C)** 100% contingency vs. 70% contingency cued fear-conditioning protocols. In the reduced contingency protocol, rats are exposed to 7 presentations of the cue light co-terminating with foot shock and 3 un-signaled foot shocks. **D)** Forward vs. backward cued fear-conditioning protocols. In the backward fear-conditioning, rats receive 10 presentations of 3.7 s cue light co-starting with 500 ms foot shock. **E)** Shock+light paired vs. un-paired cued fear-conditioning protocols. In the un-paired protocol, rats receive 8 trials of foot shocks alone and 8 cue lights alone (pseudorandomly mixed), with an addition of 2 trials of cue light co-terminating with foot shock. **F)** FPS diagram showing how cued fear, non-cued fear, and contextual fear are calculated based on the absolute acoustic startle responses (ASR).

To determine the rate of fear extinction, rats were then tested three times for cued/non-cued fear and three times for contextual fear on alternate days (unless stated otherwise).

### FPS with 100% and 70% contingency fear-conditioning protocol

Male (n=20, n=10 per group) and female SD rats (n=16, n=8 per group) were tested for ASR and FPS as described above. Rats were assigned to experimental protocols based on their pre-shock baseline ASR in order to have groups with balanced average ASR. On day 3 (fear conditioning), rats were either exposed to 10 presentations of cue light (CS) co-terminating with foot shock (100% contingency, as above), or 7 presentations of the cue light co-terminating with foot shock and 3 un-signaled foot shocks (70% contingency). On day 4, rats were tested for cued/non-cued fear, and on day 5, rats were tested for contextual fear, followed by extinction tests, as above (**Fig. 1C**).

### FPS with forward and backward fear-conditioning protocol

Male (n=28, n=14 per group) and female (n=26, n=13 per group) SD rats were assigned to each conditioning protocol and tested for ASR as above. On day 3 (fear conditioning), rats were either exposed to 10 presentations of the cue light (CS), each co-terminating with foot shock (forward conditioning, as above), or were exposed to 10 presentations of foot shocks co-starting with 3.7 s cue light (backward conditioning). On day 4, rats were tested for recall of cued/non-cued fear, and on day 5, rats were tested for contextual fear recall, followed by extinction tests, as described above (**Fig. 1D**).

### FPS with paired and un-paired fear-conditioning protocol

Male (n=34, 17 rats per group) and female (n=29, 14 rats - paired group, 15 rats - un-paired group) Wistar rats were tested for ASR as above. On day 3 (fear conditioning), one group of rats was exposed to 10 trials of cue light, each co-terminating with foot shock (shock+light paired, as above). Another group was exposed to mostly un-paired foot shocks and cue lights, where rats received 8 trials of foot shocks alone and 8 cue lights alone (pseudorandomly mixed), with an addition of 2 trials of cue light co-terminating with foot shock, in order to prevent the cue from becoming a safety signal (shock+light unpaired). On day 4, rats were tested for cued/non-cued fear, and on day 5, rats were tested for contextual fear, followed by extinction tests, as above (**Fig. 1E**).

All respective FPS protocols were tested in parallel, such that the 100% contingency protocol was directly compared with 70% contingency protocol (same for forward vs. backward conditioning protocol, and paired vs. un-paired protocol). Note, all respective control groups above (100% contingency, forward conditioning, and shock+light paired) were conditioned using the classic FPS protocol with the 3.7 s cue light co-terminating with 0.5 s foot-shock.

### FPS with contextual fear conditioning

Lastly, we used SD female (n=10) and male rats (n=10) to compare differences in contextual fear learning using FPS. In order to directly compare FPS between sexes, ASR was corrected for 100 g of body weight. Rats were tested in Plexiglas enclosures as above and ASR was measured during 30 trials of startle eliciting WNB (95 dB) on day 1 (chamber and startle habituation), and day 2 (baseline pre-shock test). On day 3 (fear conditioning, context A), rats received 10 un-signaled 0.5 s foot shocks (US; 0.5 mA). On day 4, rats were tested for contextual fear recall in context A, where post-shock ASR was measured during 30 trials of startle eliciting 95dB WNB. To determine the rate of contextual fear extinction, rats were repeatedly tested over the course of 3 days (**Fig. 1B**).

### Statistical analysis

Data are presented as mean ± SEM. Raw startle/FPS data was analyzed by a two-way repeated measures (RM) analysis of variance (ANOVA) with the factors trial type (pre-shock, post-shock, noise only, light+noise) and conditioning protocol and/or sex. In the FPS analyses, we first tested for a significant trial type effect and conditioning protocol effect comparing classic and modified fear conditioning protocols in males and females, separately. Then, we compared FPS from modified fear-conditioning protocols (70% contingency, backward, and shock+light unpaired) between males and females with trial type and sex as factors. When the F-ratio was significant, all *post-hoc* analyses were compared using Sidak’s multiple comparison test. To analyze the effect of the conditioning protocol or sex on cued, non-cued, and contextual fear as well as discrimination index and shock reactivity, data were presented as percentage change scores of ASR and were analyzed with unpaired t-tests. The percentage change of ASR was analyzed according to the following formulas (**Fig. 1F**):

Cued fear = (Light+Noise – Noise-only) / Noise-only) * 100% in context B

Noncued fear = (Noise only – Post-shock) / Post-shock) * 100% in context B

Discrimination index = (Light+Noise/Noise-only) / (Noise-only/Post-shock) in context B

Contextual fear = (Post-shock – Pre-shock) / Pre-shock) * 100% in context A

Shock reactivity for each individual rat was calculated as the average startle amplitude during each of the 10 foot-shock presentation during fear conditioning (no WNB present).

All statistical analyses were completed using GraphPad Prism version 9.4.1 (681) (GraphPad Software, Inc., San Diego, CA). *P < 0.05* was considered significant.

## Results

### 1. Female and male rats show robust FPS with 100% contingency (forward, paired) fear conditioning protocol with females demonstrating higher shock reactivity and lower fear discrimination

#### 1.1. FPS after 100% contingency protocol in females and males

Since we used the classic FPS protocol (100% contingency with forward conditioning and light and foot-shock paired) across the study to compare it with modified FPS protocols, we first compared fear expression in all female and male rats combined using the classic FPS protocol. Although males and females differ in absolute ASR (p=0.035), no significant differences were found in the ASR between males and females, when corrected for body weights (p=0.1896, **Fig. 2A**). Notably, females showed a significantly higher shock reactivity than males (p=0.0005), even more striking when corrected for body weight (p<0.0001, **Fig. 2B**). In the FPS recall in context B, two-way RM ANOVA showed a significant trial effect of noise only vs. light+noise trials (F(1,51)=36.49, p<0.0001), with *post-hoc* showing significant trial effect in both males (p<0.0001), and females (p=0.0021), suggesting an efficient cued fear acquisition and recall in both sexes. No sex effect (F(1, 51)=0.08726, p=0.7689) nor interaction between trial type (noise only vs. light+noise) and sex was observed (F(1, 51)=0.4087, p=0.5255). There was also a significant trial effect of post-shock vs. noise only trials (F(1,51)=17.46, p=0.0001) in both males (p=0.0048), and females (p=0.0148), indicating non-cued fear acquisition and recall in both sexes. No sex effect (F(1, 51)=1.925e-005, p=0.9965) nor interaction between trial type (post-shock vs. noise only) and sex was observed (F(1, 51)=0.02345, p=0.8789), (**Fig. 2C**). During contextual fear recall in context A, two-way RM ANOVA showed a significant trial effect of pre-shock vs. post-shock (F(1, 51)=12.92, p=0.0007), and a trend in sex effect (F (1, 51)=3.630, p=0.0624), while *post-hoc* shows this effect is primarily driven by males (p=0.0002) but not females (p=0.3870), suggesting that males, but not females, show a significant contextual fear recall in context A after cued fear-conditioning (**Fig. 2F**).

**Figure 2.**
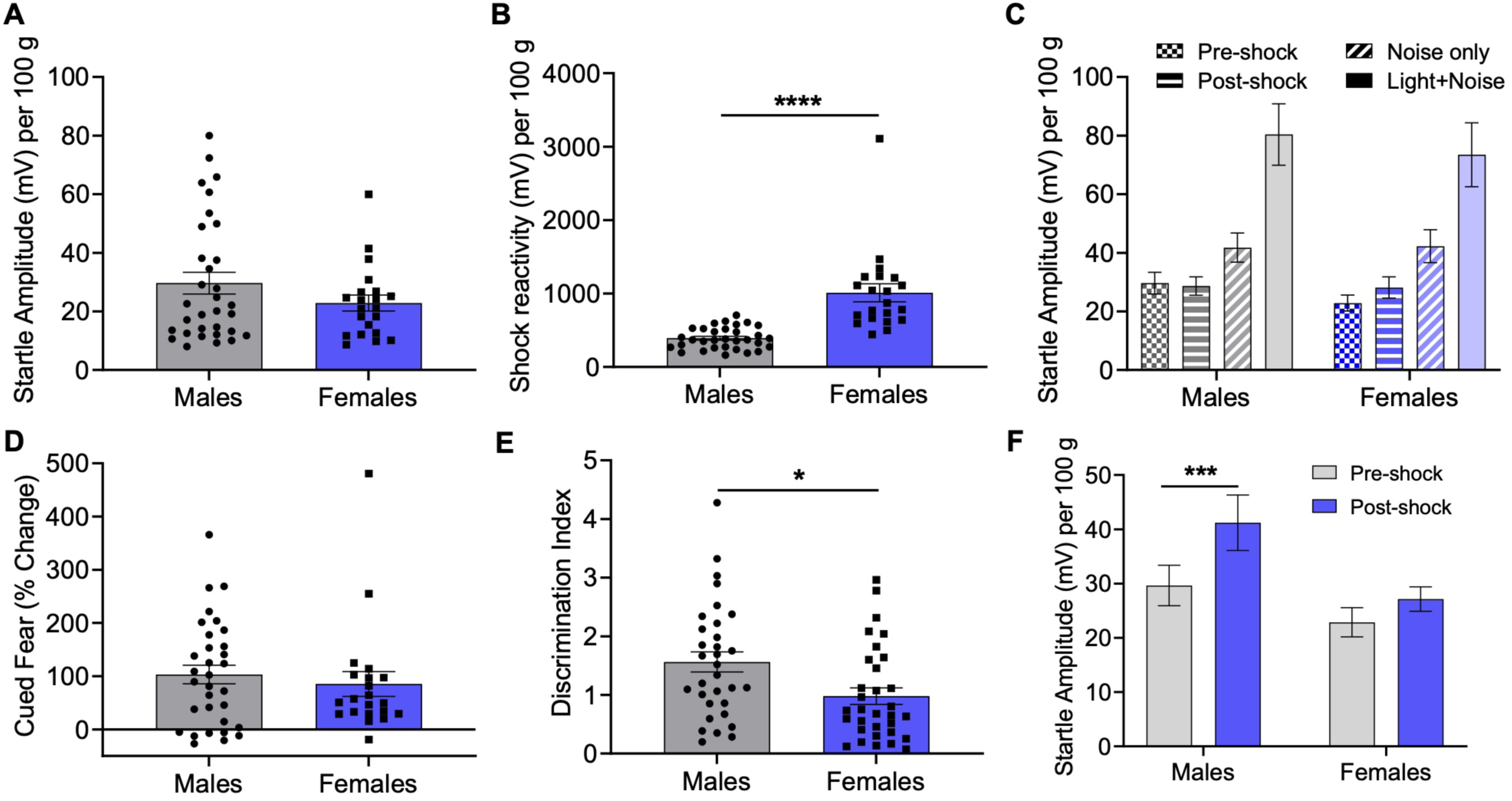
Female and male rats show robust FPS with classic fear conditioning protocol, with females showing higher reactivity to foot-shocks and reduced fear discrimination than males. **A)** Males and females show no significant differences in the startle amplitude corrected per body weight but **B)** females demonstrate significantly higher shock reactivity in comparison to males when corrected for the body weight. **C)** FPS recall in context B shows a significant trial effect between noise only vs. light + noise (p<0.0001) and a significant trial effect of post-shock vs. noise only (p=0.0001) in females and males. **D)** No significant sex differences are found in the percentage change of cued fear, but **E)** females show a significantly reduced discrimination index than males. **F)** Males, but not females, show a significant contextual fear recall in context A after cued-fear conditioning. *p<0.05, ***p<0.001, ****p<0.0001.

#### 1.2. Percentage change of fear and fear extinction following 100% contingency protocol in females and males

During the first recall tests, no significant differences were found in the percentage change of cued fear (p=0.5395, **Fig. 2D**), non-cued fear (p=0.7865, not shown), or contextual fear (p=0.4257, not shown) between males and females. However, females showed a significantly reduced discrimination index (p=0.011) than males (**Fig. 2E**), suggesting a concurrent reduction of cued and increase in non-cued fear. There was a significant effect of time on cued fear extinction (F(1.627, 82.96)=17.61, p<0.0001), but no significant effect of sex (F(1, 51)=0.007072, p=0.9333), nor significant interaction of time and sex (F(2, 102)= 0.5144, p=0.5994), suggesting that males and females extinguish cued fear in a similar rate, not shown. There was no significant effect of time on non-cued fear extinction (F(1.949, 99.41)=2.565, p=0.0834), no significant effect of sex (F(1, 51)= 0.7074, p=0.4042), nor significant interaction between sex and time (F(2, 102)=06554, p=0.5214), suggesting that non-cued fear does not follow the same rapid extinction pace as cued fear. Similarly, there was no significant effect of time F(2, 132)=0.1424, p=0.8674), no significant effect of sex (F(1, 132)=1.582, p=0.2108), nor interaction between sex and time F(2, 132)=2.459, p=0.0894) on contextual fear extinction, not shown.

These results suggest that both male and female rats show robust FPS and similar levels of cued and non-cued fear in the classic FPS, despite females showing significantly higher reactivity to foot-shocks during fear-conditioning. Notably, female rats demonstrate reduced fear discrimination in the FPS, in comparison to males.

### 2. Female and male rats show FPS with 70% and 100% contingency fear-conditioning protocols with females demonstrating significantly higher shock reactivity than males

#### 2.1. FPS after 70% vs. 100% contingency protocols in females

During fear-conditioning, no significant differences were found on shock reactivity (p=0.1457) between 70% and 100% continency protocols in females (**Fig. 3A**). During cued/non-cued fear recall in context B, two-way RM ANOVA showed a significant trial effect of noise only vs. light+noise trials (F(1, 14)=7.930, p=0.0137), but no contingency effect (F(1, 14)=0.03249, p=0.8595), nor interaction (F(1, 14)=0.9412, p=0.3484, **Fig. 3B**). There was also a significant trial effect of post-shock vs. noise only (F(1, 14)=9.663, p=0.0077); however, *post-hoc* analysis showed this is primarily driven by 70% contingency protocol (p=0.0208), and not 100% contingency (p=0.3132), **Fig. 3B.** During the contextual fear recall in context A, two-way RM ANOVA showed an overall significant trial effect (pre-shock vs. post-shock) (F(1, 14)=6.961, p=0.0195) but no contingency effect (F(1, 14)=0.08376, p=0.7765), nor interaction between trial type and contingency (F(1, 14)=0.3131, p=0.5846), not shown. Interestingly, there was a stronger trial effect during the second contextual fear recall in females (F(1, 14)=13.93, p=0.0022), but no protocol effect (F (1, 14) = 0.02380, p=0.8796), nor interaction (F (1, 14) = 0.04697, p=0.8315), suggesting that females show a more robust contextual fear expression during the second testing (**Fig. 3C**).

**Figure 3.**
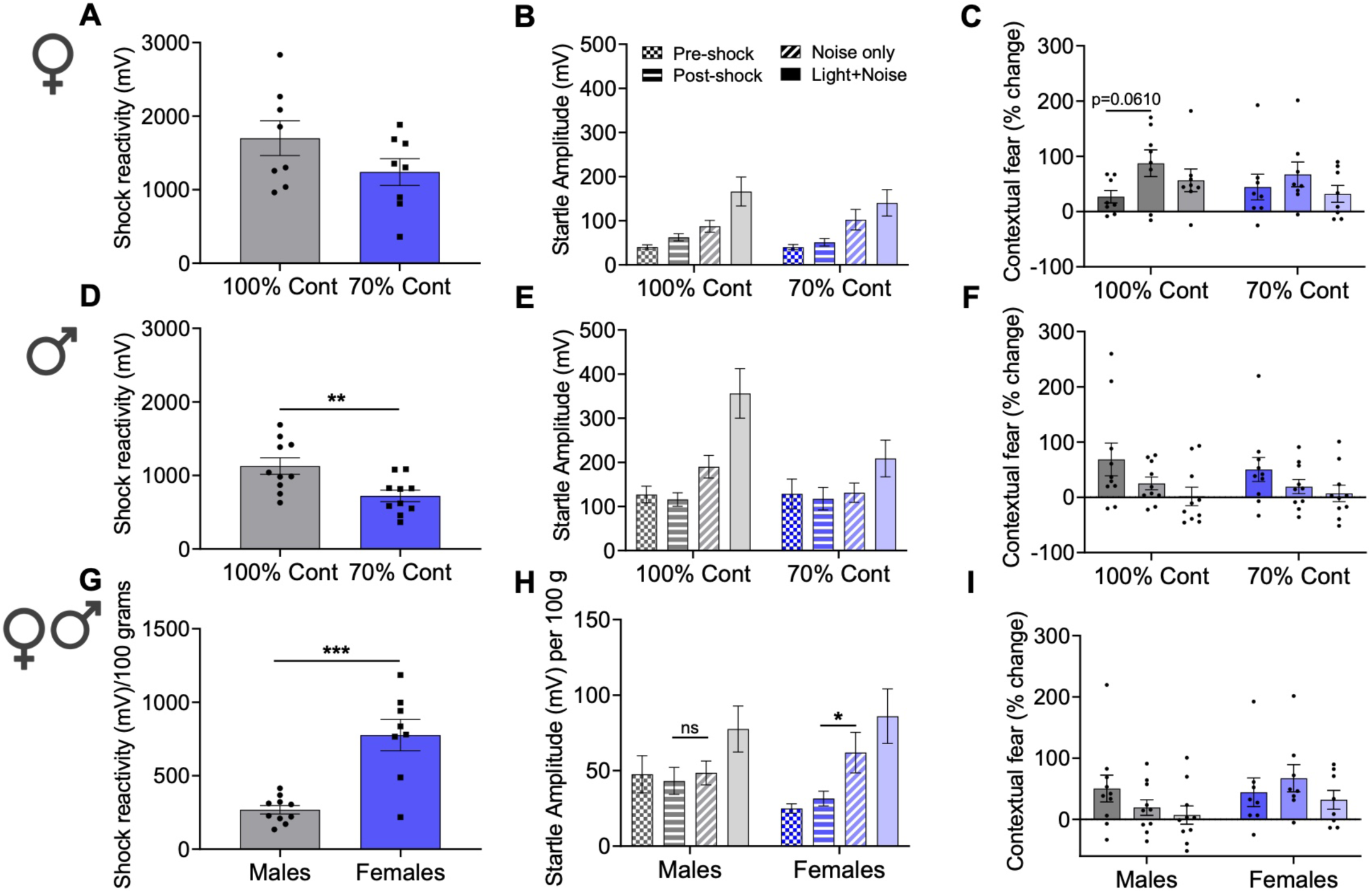
Female and male rats show FPS with 70% and 100% contingency fear-conditioning protocols, with females demonstrating significantly higher shock reactivity than males. **A)** Females show no differences in shock reactivity with 100% and 70% contingency protocols. **B)** FPS recall from 100% vs. 70% contingency fear-conditioning in females shows significant trial effect of noise only vs. light + noise (p=0.0137) and post-shock vs. noise only (p=0.0077). **C)** In females, there is a significant effect of time on contextual fear extinction (p=0.0383), but no significant effect of contingency (p=0.6745). In 100% contingency protocol, females tend to show higher contextual fear during the second test in comparison to the first test. **D)** Males demonstrate significantly higher shock reactivity with 100% vs. 70% contingency protocols. **E)** FPS recall from 100% vs. 70% contingency fear-conditioning in males shows a significant trial effect of noise only vs. light + noise (p=0.0003), and a significant effect of contingency (p=0.0447). There is also a significant trial effect of post-shock vs. noise only (p=0.0094), and a trend in the interaction between contingency and trial type (p=0.0609). **F)** Males show a significant effect of time on contextual fear extinction (p=0.0004), but no contingency effect (p=0.7945). **G)** During 70% contingency fear-conditioning, females show a significantly higher shock reactivity than males. **H)** FPS recall after 70% contingency fear-conditioning in females vs. males. Female and male rats show a significant trial effect of noise only vs. light + noise trials (p=0.0046) and a significant trial effect of post-shock vs. noise only trials (p=0.0090), and a trend in the interaction between sex and the trial type (p=0.0529). *Post-hoc* analysis show that the ASR during noise only is significantly higher than post-shock in females, but not in males. **I)** Extinction of contextual fear following 70% contingency fear-conditioning in female and male rats. **p<0.01, ***p<0.0001.

### 2.2. Percentage change of fear and fear extinction after 70% vs. 100% contingency in females

No significant differences were found in cued fear (p=0.2769), non-cued fear (p=0.4600), contextual fear (p=0.5098), or discrimination index (p=0.2221), not shown. There was a significant effect of time on cued fear extinction (F(1.391, 19.48)=5.451, p=0.0214), but no contingency effect (F (1, 14) = 0.7574, p=0.3988), nor interaction F (2, 28)= 1.044, p=0.3654), suggesting that rats extinguish cued fear at a similar rate with both contingency protocols. However, there was no effect of time on non-cued fear extinction (F(1.892, 26.49)=0.5439, p=0.5773), indicating that rats do not extinguish non-cued fear at the same rapid pace as cued fear. There was a significant effect of time on contextual fear extinction (F(1.955, 27.37)=3.712, p=0.0383), but there was no significant effect of contingency (F(1.14)=0.1840, p=0.6745), nor interaction (F (2, 28)=1.024, p=0.3724). Notably, *post-hoc* comparisons showed that females conditioned with 100% contingency tended to express higher contextual fear at the second testing in comparison with the first test (p=0.0610, **Fig. 3C**), demonstrating that, following cued fear conditioning, females show more robust contextual fear expression during the second testing.

#### 2.3. FPS after 70% vs. 100% contingency protocols in males

During fear-conditioning, shock reactivity was significantly higher in male rats conditioned with 100% contingency in comparison with 70% contingency protocol (p=0.0080, **Fig. 3D**). During cued/non-cued fear recall in context B, two-way RM ANOVA showed a significant trial effect of noise only vs. light+noise (F(1, 18)=20.43, p=0.0003), and a significant effect of contingency (F(1,18)=4.655), p=0.0447). *Post-hoc* analysis showed a significant trial effect in 100% contingency (p=0.0008), but not in 70% contingency protocol (p=0.1107), suggesting that male rats might have reduced cued fear acquisition and recall with 70% contingency protocol, **Fig. 3E.** There was a significant trial effect of post-shock vs. noise only trials (F(1, 18)=8.442, p=0.0094), and a trend in the interaction between contingency (100% vs. 70%) and trial type (post-shock vs. noise only, F(1, 18)=3.998, p=0.0609). *Post-hoc* analysis showed that 100% contingency protocol induced a significant trial effect of post-shock vs. noise only (p=0.0055), but no effect was found in 70% contingency protocol (p=0.7789, **Fig. 3E**), suggesting that 70% contingency protocol might reduce non-cued fear acquisition and recall in male rats. During the contextual fear recall in context A, two-way RM ANOVA showed a significant trial effect (pre-shock vs. post-shock) (F(1, 18)=11.76, p=0.0030), but no contingency effect ((F1, 18)=0.006079, p=0.9387), nor interaction between trial type and contingency (F(1, 18)=0.1069, p=0.7475), not shown.

### 2.4. Percentage change of fear and fear extinction after 70% vs. 100% contingency in males

No significant differences were found in cued fear (p=0.4424), non-cued fear (p=0.1820), discrimination index (p=0.7842), or contextual fear (p=0.6255) in between the protocols (not shown). There was a significant effect of time (F(1.338, 24.08)=16.59, p=0.0002), no effect of contingency protocol on cued fear extinction (F(1, 18)=0.02631, p=0.8730), nor interaction between time and contingency (F(2, 36)=1.185, p=0.3173), suggesting that males extinguish cued fear at a similar rate with both contingency protocols. Two-way RM ANOVA showed no effect of time on non-cued fear extinction (F(1, 18)=1.936, p=0.1810), no effect of contingency protocol (F1, 18)=1.936, p=0.1810), nor interaction between contingency protocol and time (F(2, 36)=2.500, p=0.0962). Finally, there was a significant effect of time on contextual fear extinction (F(1.555, 27.99)=12.37, p=0.0004), but there was no contingency effect (F1, 18)=0.06991, p=0.7945), nor interaction between time and contingency (F(2, 36)=0.5494, p=0.5820, **Fig. 3F**), indicating that males extinguish contextual fear at a similar rate with both protocols.

Together, these results show that although 70% contingency protocol reduces trial effects in the FPS in female and male rats, it does not affect percentage change of cued, non-cued, or contextual fear in comparison with 100% contingency protocol. Moreover, females and males extinguish cued fear at a similar rate with both protocols, whereas non-cued fear extinction is hindered in both protocols. However, whereas males extinguish contextual fear gradually with both protocols, females do not show a full expression of contextual fear until the second testing. Notably, males show lower shock reactivity with 70% protocol in comparison to 100% contingency protocol, an effect not observed in females.

### 2.5. FPS after 70% contingency protocol in females vs. males

During fear-conditioning with 70% contingency protocol, females showed significantly higher shock reactivity (p=0.0122) than males, even more striking when corrected for body weight (p=0.0001, **Fig. 3G**). To directly compare the FPS between males and females, ASR was corrected for body weight. Two-way RM ANOVA showed a significant trial effect of noise only vs. light+noise trials (F(1, 16)=10.82, p=0.0046), but no sex effect (F(1, 16)=0.3750, p=0.4890), nor interaction (F(1, 16)=0.09456, p=0.7624, **Fig. 3H**). Two-way RM ANOVA showed a significant trial effect of post-shock vs. noise only (F(1, 16)=8.844, p=0.0090), no sex effect (F(1, 16)=0.005588, p=0.9413), and a trend in the interaction between sex and post-shock vs. noise only trials (F(1, 16)=4.370, p=0.0529), **Fig. 3H.** *Post-hoc* analysis showed that the ASR during noise only trial is significantly higher than post-shock in females (p=0.0339), but not in males (p=0.6830), indicating that 70% contingency might produce non-cued fear primarily in females. For the contextual fear recall in context A, two-way RM ANOVA showed a significant trial effect (pre-shock vs. post-shock) (F(1, 16)=7.209, p=0.0163), a trend in sex effect (F(1, 16)=3.938, p=0.0646), but no interaction between trial type and sex (F(1, 16)=0.5111, p=0.4849). Here, *post-hoc* analysis showed that ASR is significantly higher during post-shock vs. pre-shock in males (p=0.0424), but not in females (p=0.3678). Interestingly, for the second contextual fear recall, results also showed a trial effect (F(1, 16)=6.479, p=0.0216), but no effect of sex (F(1, 16)=1.214, p=0.2869), nor interaction between trial and sex (F(1, 16)=2.88, p=0.1092), not shown. *Post-hoc* analysis showed that ASR is significantly higher during post-shock vs. pre-shock in females (p=0.0233), but not in males (p=0.7821), suggesting that females, but not males, show higher contextual fear expression during the second testing (**Fig. 3I**).

### 2.6. Percentage change of fear and fear extinction after 70% contingency protocol in females vs. males

No significant differences were found in a percentage change of cued fear (p=0.6680), non-cued fear (p=0.1311), discrimination index (p=0.3041), or contextual fear expression (p=0.8517) between males and females (not shown). However, during the second contextual fear testing females tended to show higher contextual fear than males (p=0.0674), not shown. There was a significant effect of time on cued fear extinction (F(1, 950, 31.20)=6.2664, p=0.0055), but no sex effect (F(1, 16)=0.2176, p=0.6471) nor interaction (F(2, 32)=0.1451, p=0.8655). There was no effect of time on non-cued fear extinction (F(1.966, 31.46)= 0.1213, p=0.8830), no sex effect (F(1, 16)=0.3559, p=0.5591), and no interaction between sex and time (F(2, 32)= 1.678, p=0.2028). There was a trend in time effect (F(1.843, 29.48)=2.546, p=0.0993), no sex effect (F(1, 16)=1.073, p=0.3156), and no interaction between sex and time during contextual fear extinction (F(2, 32)=2.066, p=0.1433, **Fig. 3I**).

Overall, these results show that although 70% contingency protocol produced stronger trial effect pointing toward more robust contextual fear in males and more robust non-cued fear in females, no differences in percentage change of cued, non-cued fear, or contextual fear were found between sexes during the first recall tests. However, male and female vary in contextual fear expression over time, such as following 100% contingency protocol, females tend to show higher contextual fear than males during the second testing. In agreement with other protocols, females show significantly higher reactivity to foot-shocks during fear conditioning.

### 3. Female and male rats show reduced FPS after backward in comparison to forward fear conditioning, demonstrated as reduced cued fear and increased contextual fear

### 3.1. FPS after forward vs. backward conditioning protocols in females

During fear-conditioning, backward conditioning protocol reduced shock reactivity in comparison with forward conditioning in females (p=0.0137, **Fig. 4A**). During cued/non-cued fear recall in context B, two-way RM ANOVA showed a significant trial effect of light+noise vs. noise only (F(1, 24)=6.984, p=0.0143) and no significant protocol effect (F(1, 24)=1.640, p=0.2126). However, there was a significant interaction between conditioning protocol (forward vs. backward) and this trial type (F(1, 24)=5.408, p=0.0288), **Fig. 4B.** *Post-hoc* analysis showed that ASR is significantly higher during light+noise trial vs. noise only trial with forward protocol (p=0.0360) but not with backward protocol (p=0.9692) in females. There was also a significant trial effect of post-shock vs. noise only (F(1, 24)=5.140, p=0.0327), but no protocol effect (F(1, 24)=0.001432, p=0.9701), nor interaction (F(1, 24)=0.6999, p=0.4111, **Fig. 4B**). During contextual fear recall in context A, two-way RM ANOVA showed a significant trial effect (pre-shock vs. post-shock, (F1, 24)=12.13, p=0.0019)), a significant effect of protocol (F(1, 24)=4.874, p=0.0371), and a significant interaction between protocol and trial type (F(1, 24)=8.379, p=0.0080). *Post-hoc* analysis showed that ASR is significantly higher during post-shock vs. pre-shock with backward conditioning protocol (p=0.0003), but no significant differences were found with forward conditioning protocol (p=0.8985), suggesting that females show greater contextual fear with backward conditioning protocol, as oppose to forward conditioning during the first test, not shown.

**Figure 4.**
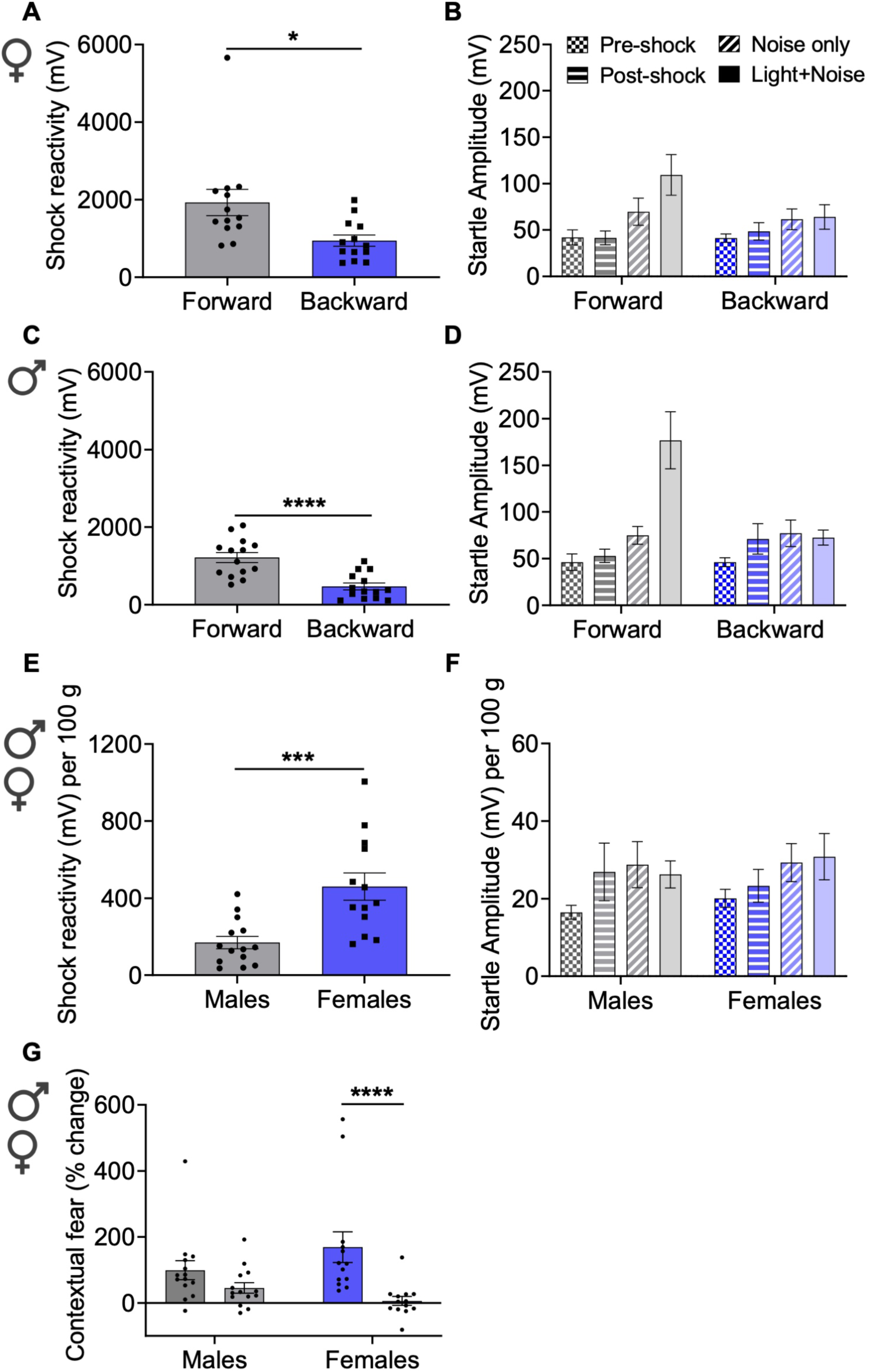
Female and male rats show reduced shock reactivity during backward vs. forward fear-conditioning. **A)** Females show significantly reduced shock reactivity with backward fear-conditioning in comparison with forward conditioning. **B)** FPS recall after forward vs. backward fear-conditioning in females shows a significant trial effect (p=0.0143) of light + noise vs. noise only and a significant interaction between conditioning protocol and the trial type (p=0.0288). There is also a significant trial effect of post-shock vs. noise only (p=0.0327) but no protocol effect (p=0.9701). **C)** Males demonstrate reduced shock reactivity with backward vs. forward conditioning. **D)** FPS recall after forward vs. backward fear-conditioning in males shows a trial effect of noise only vs. light + noise (p=0.0019), a significant effect of protocol (p=0.0227), and a significant interaction (p=0.0008). There is also a trend in trial effect of post-shock vs. noise only (p=0.0646), but no effect of conditioning protocol (p=0.5264). **E)** During backward fear conditioning, females show a significantly higher shock reactivity than males. **F)** Females and males show similar FPS recall following backward fear-conditioning. **G)** There is a significant effect of time (p<0.0001), no sex effect (p=0.6518), but a significant interaction of sex and time (p=0.0259) on the contextual fear extinction, with a significant difference between contextual fear tests 1 and 2 in females but not males. *p<0.05, ***p<0.01, ****p<0.0001.

### 3.2. Percentage change of fear and fear extinction after forward vs. backward conditioning in females

Backward fear conditioning tended to reduce cued fear in comparison with forward conditioning (p=0.0873, **Fig. 5A**), but no significant differences were found in non-cued fear (p=0.3222, not shown), or discrimination index (p=0.8764, **Fig. 5B**). However, the percentage change of contextual fear was significantly higher with backward conditioning protocol in comparison with forward fear conditioning (p=0.0124, **Fig. 5C**) in female rats. Two-way RM ANOVA showed no significant effect of time (F(1.848, 44.35)=2.036, p=0.1458), but a significant effect of protocol on cued fear extinction (F(1, 24)=5.930, p=0.0227), and no significant interaction between time and protocol (F(2, 48)=1.460, p=0.2424). However, *post-hoc* analysis showed no significant differences between protocols in the cued fear tests (test 1: p=0.2486, test 2: p=0.1277, test 3: p=0.9769), not shown. There was a trend of time effect on non-cued fear extinction (F(1.609, 38.61)=2.766, p=0.0859), no significant effect of protocol (F(1, 24)=0.4375, p=0.5146), nor interaction between time and protocol (F(2, 48)=1.034, p=0.3634), not shown. There was a significant effect of time (F(1, 24)=10.57, p=0.0034), a significant effect of protocol (F(1, 24)=7.844, p=0.0099), and a significant interaction between conditioning protocols and time on contextual fear extinction (F(1, 24)=4.979, p=0.0353), and a *post-hoc* analysis showed a significantly higher contextual fear following backward, as opposed to forward conditioning, during the first contextual test in females (p=0.0020, not shown), confirming the results above (**Fig. 5C**).

**Figure 5.**
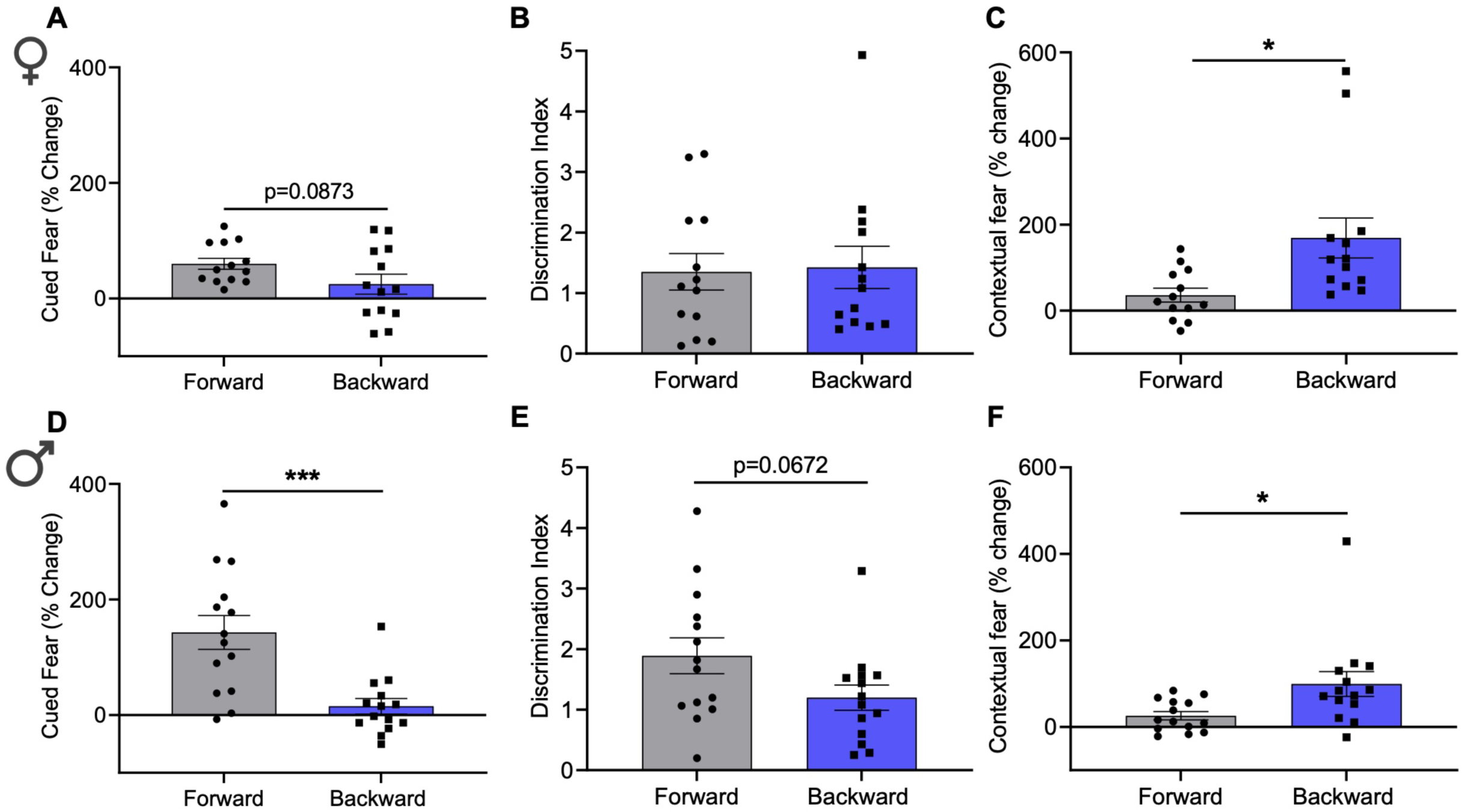
Male rats show reduced cued fear, whereas both sexes show enhanced contextual fear after backward vs. forward fear-conditioning. **A)** Females tend to show reduced cued fear **B)** but show no changes in the discrimination index after backward vs. forward fear-conditioning. **C)** Females demonstrate enhanced contextual fear after backward vs. forward fear-conditioning. **D)** Males show significantly reduced percentage change of cued fear and **E)** tend to reduce the discrimination index after backward vs. forward fear-conditioning. **F)** Males show significantly higher percentage change of contextual fear after backward vs. forward fear-conditioning. *p<0.05, ***p<0.001.

#### 3.3. FPS after forward vs. backward conditioning protocols in males

During fear-conditioning, backward conditioned males showed significantly lower shock reactivity (p<0.0001, **Fig. 4C**). Two-way ANOVA showed a significant trial effect of noise only vs. light+noise (F(1, 26)=11.93, p=0.0019), a significant effect of conditioning protocol (F(1, 26)=5.869, p=0.0227), and a significant interaction between protocol (forward vs. backward) and these trials (F(1, 26)=14.31, p=0.0008). *Post-hoc* analysis showed that ASR is significantly higher during light+noise trial vs. noise only trial with forward conditioning (p<0.0001), but no significant differences were found with backward conditioning protocol (p=0.9667), **Fig. 4D.** Results showed a trend effect of post-shock vs. noise only trials (F(1, 26)=3.725, p=0.0646), but no effect of conditioning protocol (F(1, 26)=0.4123, p=0.5264), nor interaction (F(1, 26)=1.189, p=0.2855, **Fig. 4D**). During contextual fear recall in context A, two-way RM ANOVA showed a trial effect (pre-shock vs. post-shock, (F(1, 26)=11.39, p=0.0023)), no significant effect of conditioning protocol (F(1, 26)=1.200, p=0.2833), and a trend in the interaction between trial and conditioning protocol (F(1, 26)=3.648, p=0.0672). *Post-hoc* analysis showed that ASR is significantly higher during post-shock vs. pre-shock with backward conditioning (p=0.0019), but not with forward conditioning (p=0.5237) in males, **Fig. 4D**.

#### 3.4. Percentage change of fear and fear extinction after forward vs. backward conditioning protocols in males

Forward conditioned rats showed significantly higher cued fear in comparison with backward conditioned rats (p=0.0005, **Fig. 5D**), but no significant differences were found in non-cued fear (p=0.1804), not shown. However, forward conditioned rats tended to have higher fear discrimination index than backward conditioned rats (p=0.0672, **Fig. 5E**). Similarly to females, backward conditioned rats showed significantly higher percentage change of contextual fear than forward conditioned rats (p=0.0225, **Fig. 5F**). There was a significant effect of time (F(1.861, 48.38)=6.214, p=0.0048), a significant effect of conditioning protocol (F(1, 26)=13.53, p=0.0011), and a significant interaction between protocol (forward vs. backward) and time on cued fear extinction (F(2, 52)=6.641, p=0.0027), with *post-hoc* analysis showing significantly higher cued fear during the first recall test after forward conditioning (p=0.0026), confirming the results above. No significant effect of time was found on non-cued fear extinction (F(1.984, 51.59)=1.138, p=0.3280), and no significant protocol effect (F(1, 26)=0.3502, p=0.5591), nor interaction (F(2, 52)=1.202, p=0.3087), not shown. However, although there was no significant time effect (F(1.857, 48.29)=2.385) on contextual fear extinction, there was a trend effect of conditioning protocol (F(1, 26)=3.877, p=0.0597), and a significant interaction between conditioning protocol and time (F(2, 52)=4.450, p=0.0164). Again, *post-hoc* analysis showed that backward conditioned rats tend to have higher contextual fear than forward conditioned rats during the first recall test (p=0.0802), confirming the results above.

Together these results indicate that, in comparison to forward conditioning, backward conditioning tends to reduce cued fear in female rats and robustly reduces cued fear in male rats but it does not affect non-cued fear in either sex. Furthermore, in both sexes backward conditioning significantly reduces reactivity to foot shocks, and increases contextual fear expression in comparison with forward conditioning.

#### 3.5. FPS after backward fear-conditioning in females vs. males

Females showed a significantly higher shock reactivity than males (p=0.0094), even more striking when corrected for body weight (p=0.0007, **Fig. 4E**). Two-way RM ANOVA showed no trial effect of light+noise and noise only trials (F(1, 25)=0.02062, p=0.8870), no sex effect (F(1, 25)=0.1585, p=0.6939), and no interaction between trial and sex (F(1, 25)=0.3341, p=0.5684), **Fig. 4F.** There was no significant trial effect of post-shock vs. noise only trials (F(1, 25)=1.212, p=0.2813), no sex effect (F(1, 25)=0.04193, p=0.8394), and no interaction (F(1, 25)=0.3346, p=0.5681, **Fig. 4F**). During the contextual fear recall in context A, two-way RM ANOVA showed a significant trial type effect (pre-shock vs. post-shock, (F(1, 25)=18.52, p=0.0002), but there was no sex effect (F(1, 25)=2.333, p=0.1392), nor interaction between trial type and sex (F(1, 25)=1.848, p=0.1862), not shown.

#### 3.6. Percentage change of fear and fear extinction after backward fear-conditioning in females vs. males

No significant differences were found between males and females in the percentage change of cued fear (p=0.6635), non-cued fear (p=0.4876), discrimination index (p=0.6732), or contextual fear (p=0.2050), not shown.No significant effect of time (F(1.831, 45.78)=0.5998, p=0.5390), or sex (F(1, 25)=0.01312, p=0.9097), nor significant interaction between sex and time (F(2, 50)= 0.9319, p=0.4005) on non-cued fear extinction. There was a significant effect of time (F(1, 25)=22.21, p<0.0001), but no sex effect (F(1, 25)=0.2087, p=0.6518) on contextual fear extinction with backward conditioning. However, there was a significant interaction of sex and time (F(1, 25)=5.612, p=0.0259), and *post-hoc* showed a significant difference between contextual fear tests 1 and 2 in females (p<0.0001), but not males (p=0.1966), **Fig. 4G.**

These results indicate that males and females show similar FPS, cued, non-cued, and contextual fear expression when conditioned with backward protocol, despite vast differences in reactivity to foot-shocks.

### 4. Female and male rats show no FPS after unpaired fear conditioning with female rats demonstrating higher shock reactivity, significantly higher ASR overall, and higher contextual fear

#### 4.1. FPS after paired vs. unpaired fear-conditioning protocols in females

During fear-conditioning, there were no significant differences in shock reactivity between the protocols (p=0.9822), not shown. During cued/non-cued fear recall in context B, two-way ANOVA showed a significant trial effect of noise only vs. light+noise (F(1, 27)=7.266, p=0.0119), no protocol effect (F(1, 27)=1.464, p=0.2367), and a significant interaction between conditioning protocols (paired vs. unpaired) and these trials (F(1, 27)=4.399, p=0.0455). *Post-hoc* analysis showed that the ASR is significantly higher during light+noise than during noise only trial in paired conditioned rats (p=0.0050), but not in unpaired rats (p=0.8913), Fig. 6A. Results showed a significant trial effect of post-shock vs. noise only (F(1, 27)=9.262, p=0.0052), but no protocol effect (F(1, 27)=1.573, p=0.2206), nor interaction between this trial type and protocol (F(1, 27)=2.460, p=0.1284, Fig. 6A). During contextual fear recall in context A, two-way RM ANOVA showed a significant trial effect (pre-shock vs. post-shock, F(1, 27)=8.975, p=0.0058), but no protocol effect (F(1, 27)=1.533, p=0.2264), nor interaction (F(1, 27)=2.822, p=0.1045). *Post-hoc* analysis showed that ASR is significantly higher during post-shock in unpaired conditioned rats (p=0.0046), but not in paired conditioned rats (p=0.6011), indicating higher contextual fear recall with unpaired vs. paired protocol in female rats, Fig. 6D.

Female rats also seem to have an overall higher ASR at the beginning of FPS recall session in context B (post-shock trials) following unpaired protocol. Indeed, two-way RM ANOVA showed a significant trial effect (pre-shock vs. post-shock in context B, F(1, 27)=13.09, p=0.0012), no protocol effect (F(1, 27)=2.292, p=0.1417), and a significant interaction between these trials and protocol (F(1, 27)=6.982, P=0.0135). *Post-hoc* analysis showed a significant difference in post-shock ASR between paired and unpaired protocols (p=0.0228), and a significant difference between pre-and post-shock ASR in unpaired (p=0.0002), but not paired protocol in female rats (p=0.7531), **Fig. 6A**.

**Figure 6.**
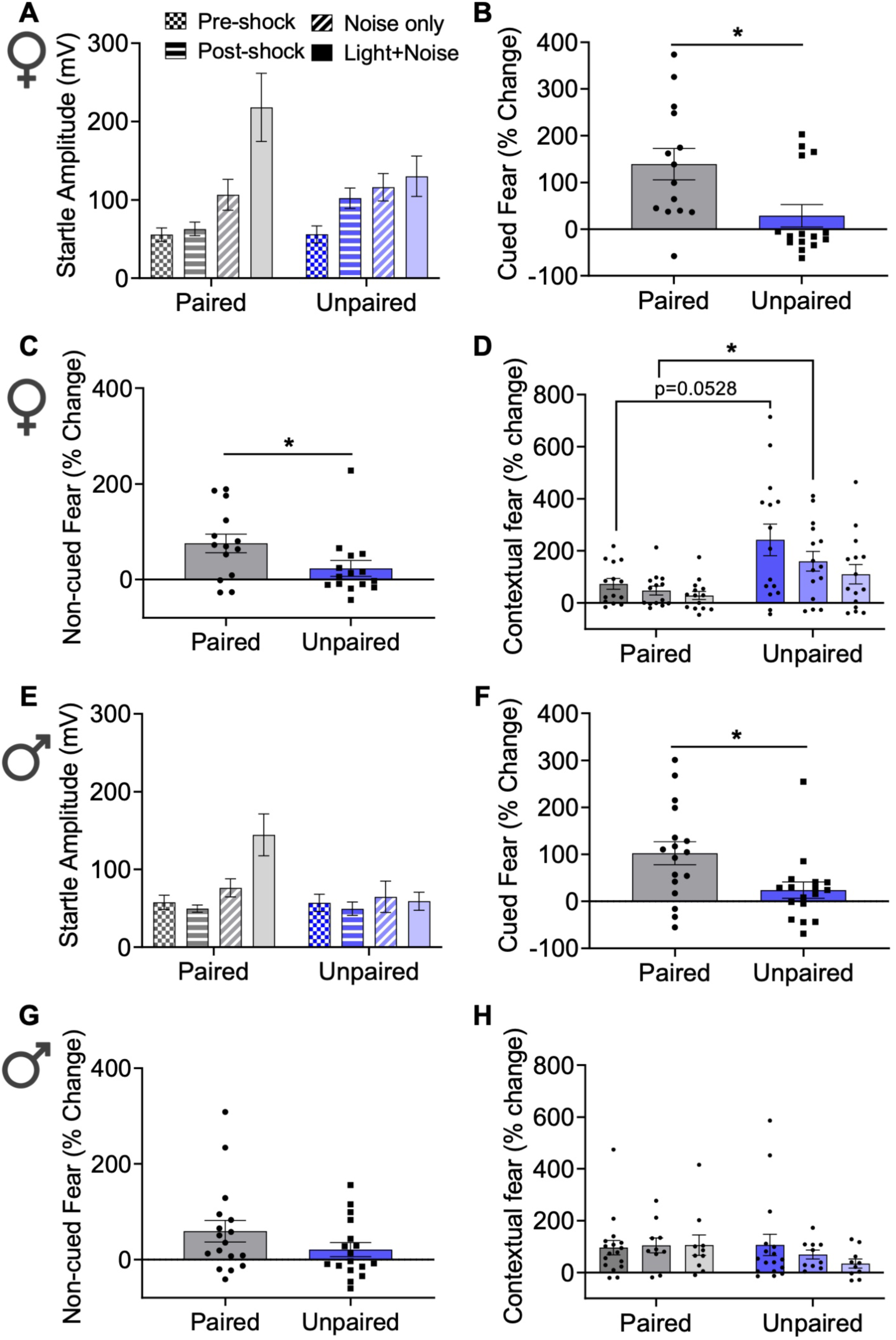
Both sexes show reduced cued fear after unpaired vs. paired fear-conditioning with females also showing reduced non-cued fear and higher contextual fear. **A)** FPS recall after paired vs. un-paired fear-conditioning in females shows a significant trial effect of noise only vs. light + noise (p=0.0119) and a significant interaction between protocols and the trials (p=0.0455). There is a significant trial effect of post-shock vs. noise only (p=0.0052), but no protocol effect (p=0.2206). Lastly, there is also a significant trial effect in pre-shock vs. post-shock in context B (p=0.0012) and a significant interaction between these trial types and protocol (P=0.0135), with *post-hoc* showing a significantly higher post-shock ASR in unpaired vs. paired protocol (p=0.0228). **B)** Females show reduced cued fear, **C)** and reduced non-cued fear following unpaired vs. paired fear-conditioning protocol. **D)** Females show a significant effect of time (p=0.0012) and a significant effect of protocol (p=0.0128) on the contextual fear extinction, with *post-hoc* indicating higher contextual fear during the first and second test following un-paired vs. paired protocol. **E)** FPS recall after paired vs. unpaired fear-conditioning in males shows a significant trial effect (p=0.0109), a significant effect of conditioning protocol (p=0.0498), and a significant interaction (p=0.0032). There is also a significant trial effect of post-shock vs. noise only trials (p=0.0168), with no protocol effect (p=0.7153). **F)** Cued fear is reduced in male rats subjected to unpaired vs. paired fear-conditioning protocol, **G)** with no significant differences in non-cued fear. **H)** Males show no significant effect of time (p=0.2592) or protocol (p=0.2592) on the contextual fear extinction.

#### 4.2. Percentage change of fear and fear extinction after paired vs. unpaired fear-conditioning protocols in females

Unpaired protocol reduced cued fear (p=0.0117, **Fig. 6B**) and non-cued fear (p=0.0495, **Fig. 6C**) in comparison to paired protocol. No differences were found in fear discrimination (p=0.3073, not shown). Yet, unpaired protocol induced a significantly higher contextual fear expression in comparison with paired conditioning in female rats (unpaired t-test, p=0.0170), **Fig. 6D.** There was a significant effect of time (F(2, 68)=5.118, p=0.0085), a significant effect of conditioning protocol (F(1, 68)=4.354, p=0.0407), and a trend in the interaction between conditioning protocol (paired vs. unpaired) and time on cued fear extinction (F(2, 68)=2.896, p=0.0621). *Post-hoc* analysis showed that cued fear is significantly higher in paired conditioned rats than in unpaired rats during the first recall test (p=0.0058), but no differences were found during the second (p=0.2807) or the third test (p=0.9106), not shown. No significant effect of time (F(2, 68)=0.3424, p=0.7133), no effect of protocol (paired vs. unpaired, F(1, 68)= 2.665, p=0.1072), nor interaction between time and protocol (F(2, 68)=1.893, p=0.1584) was found on non-cued fear extinction, not shown. However, there was a significant effect of time (F(1.628, 43.96)=8.867, p=0.0012), and a significant effect of protocol (F(1, 27)=7.114, p=0.0128), but no interaction (F(2, 54)=2.199, p=0.1207) on contextual fear extinction. *Post-hoc* analysis indicated that contextual fear is higher during the first (p=0.0528) and second test (p=0.0420) in unpaired conditioned rats in comparison with paired protocol. No significant differences were found during the third test in between the protocols (p=0.1635), **Fig. 6D**.

These results suggest that although female rats conditioned with unpaired vs. paired protocol show significantly reduced cued and non-cued fear expression, they demonstrate a significantly higher ASR during post-shock trials (in context B), in comparison to paired protocol. Furthermore, females show significantly higher levels of contextual fear in unpaired in comparison to paired protocol.

#### 4.3. FPS after paired vs. unpaired conditioning protocols in males

No significant differences were found in shock reactivity (p=0.9381) in between the protocols, not shown. Two-way RM ANOVA showed a significant trial effect (F(1, 32)=7.307, p=0.0109), a significant effect of conditioning protocol (F(1, 32)=4.157, p=0.0498), and a significant interaction between conditioning protocol (paired vs. unpaired) and noise only vs. light+noise trials (F(1, 32)=10.19, p=0.0032). *Post-hoc* analysis showed that ASR is significantly higher during light+noise in comparison to noise only trial in rats conditioned with paired protocol (p=0.0004), but not with unpaired protocol (p=0.9280), **Fig. 6E.** There was also a significant trial effect of post-shock vs. noise only trials (F(1, 32)=6.364, p=0.0168), with no protocol effect (F(1, 32)=0.1354, p=0.7153), and no interaction (F(1, 32)=0.4705, p=0.4977, **Fig. 6E**). During contextual fear recall in context A, two-way RM ANOVA showed a significant trial effect (pre-shock vs. post-shock, F(1, 32)=23.60, p<0.0001), but no protocol effect (F(1, 32)=0.05666, p=0.8134), nor interaction (F(1, 32)=0.1982, p=0.6592). *Post-hoc* analysis showed that ASR is significantly higher during post-shock vs. pre-shock in paired conditioned males (p=0.0014), as well as in unpaired conditioned males (p=0.0076), suggesting that both protocols produce contextual fear in males, not shown.

To assess potential changes in ASR post-conditioning during FPS recall, similarly to females, we compared pre-shock ASR with post-shock ASR in context B. In contrast to females, two-way RM ANOVA did not show any trial effect F (1, 18) = 2.197, p=0.1556), no protocol effect (F (1, 18) = 0.1273, p=0.7254), nor interaction between trial and protocol in male rats (F (1, 18) = 0.2887, p=0.5976), not shown.

#### 4.4. Percentage change of fear and fear extinction after paired vs. unpaired protocols in males

Cued fear was significantly higher in rats conditioned with paired vs. unpaired protocol (p=0.0135, **Fig. 6F**). However, no significant differences were found in non-cued fear (p=0.1624, **Fig. 6G**), discrimination index (p=0.6748, not shown), or contextual fear (p=0.8292, not shown). There was a significant effect of time (F(2, 82)=7.681, p=0.0009), no effect of conditioning protocol (F(1, 82)=0.7841, p=0.3785), and a significant interaction between conditioning protocol and time on cued fear extinction (F(2, 82)=5.275, p=0.0070). *Post-hoc* analysis showed that cued fear is significantly higher in paired conditioned rats in comparison with unpaired conditioned rats during the first cued fear recall test (p=0.0039), but no significant differences were found during the second (p>0.9999) or the third test (p=0.4957), not shown. No significant effect of time (F(2, 82)=0.07485, p=0.9280) was found on non-cued fear extinction. However, there was a significant effect of conditioning protocol (paired vs. unpaired, F(1, 82)=6.873, p=0.0104). Still, no significant differences were found in the non-cued fear recall test in between protocols (test 1: p=0.4464, test 2: p=0.1639, test 3: p=0.4483), not shown. No significant effect of time (F(2, 68)=1.295, p=0.2592), conditioning protocol (F(1, 68)=1.295, p=0.2592), nor interaction between protocol and time was found in contextual fear extinction (F(2, 68)=0.8066, p=0.4506, **Fig. 6H**)

Together, the results show that in comparison to paired conditioning protocol, unpaired protocol reduces cued fear and non-cued fear but does not affect contextual fear in male rats. In addition, in contrast to females, male rats do not demonstrate a significantly higher ASR overall following unpaired conditioning, in comparison to paired conditioning protocol.

#### 4.5. FPS after unpaired conditioning protocol in females vs. males

During fear-conditioning with unpaired protocol, shock reactivity was significantly higher in females (p=0.0105) even more striking when corrected for body weight (p=0.0026, **Fig. 7A**). Two-way RM ANOVA showed no significant trial effect in light+noise vs. noise only (F(1, 30)=0.2004, p=0.6576). However, there was a significant sex effect on light+noise vs. noise only trials (F(1, 30)=12.63, p=0.0013). *Post-hoc* analysis showed significantly higher ASR during light+noise (p=0.0022) and noise only trials (p=0.0141) in females in comparison to males, **Fig. 7B**. There was a trend in trial effect of post-shock vs. noise only trials (F(1, 30)= 3.056, p=0.0907), and a significant sex effect of post-shock vs. noise only trials (F(1, 30)= 14.03, p=0.0008). *Post-hoc* analysis showed a significantly higher ASR during post-shock (p=0.0030) and noise only (p= 0.0012) in females than males (**Fig. 7B**).

**Figure 7.**
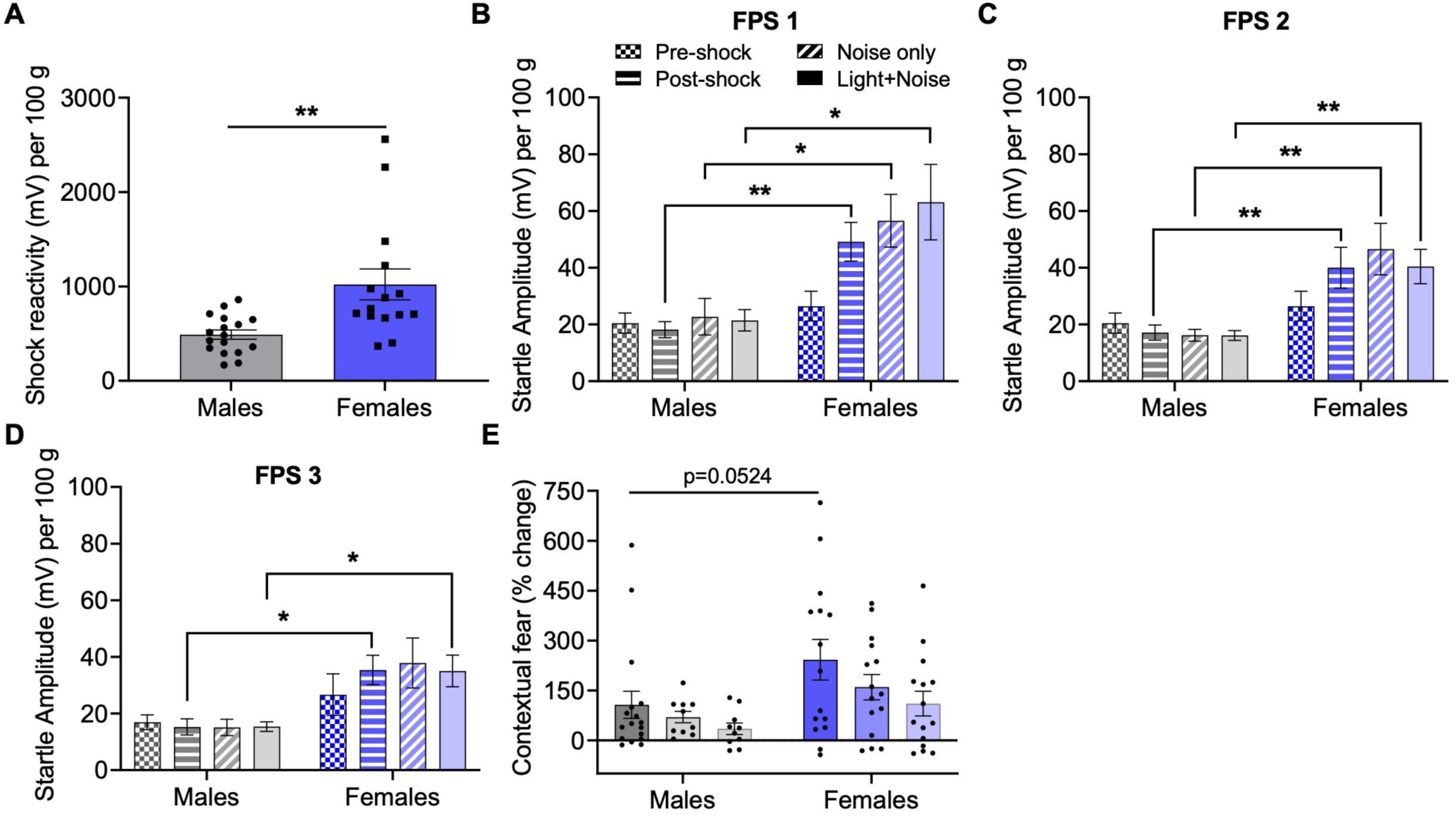
Females show increased shock reactivity, enhanced contextual fear, and higher ASR overall following unpaired fear-conditioning in comparison to males. **A)** Females show significantly higher shock reactivity than males. **B)** During the first FPS recall after unpaired fear conditioning, there is a significant trial effect overall (p=0.0073), a significant protocol effect (p=0.0008), and a significant trial and protocol interaction (p=0.0053). *Post-hoc* analysis shows that although females and males do not differ in pre-shock startle (p=0.8262), females have higher ASR at post-shock, noise only, and light + noise trials than males. **C)** During the second FPS test, there is no significant trial effect (p=0.4727) but a significant effect of sex (p<0.0001). Although males and females do not differ in pre-shock startle (p=0.8730), females show higher post-shock, noise only, and light + noise ASR than males. **D)** During the third FPS test, there was no significant trial effect (p=0.6137) but a significant effect of sex (p=0.0020), with no sex differences in pre-shock ASR (p=0.6708), but significantly higher ASR in females than males at post-shock and light + noise. E) During contextual fear extinction, there is a trend effect of time (p=0.0582), a significant effect of sex (p=0.0062), with females tending to show a higher contextual fear than males during the first test.

The results above suggest that during FPS recall test in context B, females have higher ASR overall than males, regardless of trial type. Hence, we also performed two-way RM ANOVA with sex and all trial types to determine if this was the case. Results showed a significant trial effect (F (2.033, 61.00) = 5.301, p=0.0073), a significant protocol effect (F (1, 30) = 14.02, p=0.0008), and a significant trial and protocol interaction (F (3, 90) = 4.520, p=0.0053). *Post-hoc* analysis showed that although females and males did not differ in pre-shock ASR (p=0.8262), females had higher ASR at post-shock (p=0.0020), noise only (p=0.0240), and light+noise trials (p=0.0322) (**Fig. 7B**). To determine if the higher ASR in females following unpaired protocol persisted over repeated testing, we also performed two-way RM ANOVA on data from subsequent FPS tests 2 and 3. Results from FPS 2 showed no significant trial effect (F(3, 120)=0.8435, p=0.4727), a significant effect of sex (F(1, 120)=34.00, p<0.0001), and no trial and sex interaction F(3, 120) = 2.126, p=0.1005). *Post-hoc* analysis showed that although there was no sex difference in pre-shock ASR (p=0.8730), females again showed higher ASR at post-shock (p=0.0072), noise only (p=0.0002), and light+noise (p=0.0038) than males (**Fig. 7C**). During FPS 3 results again showed no significant trial effect (F(2.532, 40.52)=0.5632, p=0.6137), a significant effect of sex (F(1, 16)=13.67, p=0.0020), and no trial and sex interaction (F(3, 48)=1.115, p=0.3522). *Post-hoc* analysis showed again no sex differences in pre-shock ASR (p=0.6708), but females showed higher ASR at post-shock (p=0.0241), and light+noise (p=0.0364) than males, although there were no sex differences in noise only (p=0.1460, **Fig. 7D**).

During contextual fear recall in context A, two-way RM ANOVA showed a significant trial effect (pre-shock vs. post-shock, F(1, 30)=10.71, p=0.0027), a trend in sex effect (F(1, 30)=3.818, p=0.0601), and a trend in the interaction between trial type and sex (F(1, 30)=3.534, p=0.0699). *Post-hoc* analysis showed that ASR is significantly higher during post-shock in context A in females (p=0.0027) but not in males (p=0.5336). Furthermore, although there was no sex difference in pre-shock (p=0.9085), females had higher ASR at post-shock (p=0.0181), not shown.

#### 4.6. Percentage change of fear and fear extinction after paired vs. unpaired conditioning protocols in females vs. males

No significant differences between males and females were found in cued fear (p=0.8640), non-cued fear (p=0.9213), or discrimination index (p=0.5644) following unpaired conditioning, not shown. However, contextual fear tended to be higher in females than in males (unpaired t-test, p=0.0698). There was no significant time effect (F(2, 76)=0.9035, p=0.4095), no sex effect (F(1, 76)=0.03580, p=0.8504), nor interaction between time and sex (F(2, 76)=0.1040, p=0.9014) on cued fear extinction, not shown. Similarly, there was no significant time effect (F(2, 76)= 0.9804, p=0.3798), no sex effect (F(1, 76)=0.002781, p=0.9581), nor significant interaction between time and sex (F(2, 76)=0.1149, p=0.8917) on non-cued fear extinction, not shown. Notably, there was a trend of time (2, 76)=2.953, p=0.0582), a significant effect of sex on contextual fear extinction (F(1, 76)=7.940, p=0.0062), but no significant interaction between time and sex (2, 76)=0.2787, p=0.7575). *Post-hoc* analysis showed that females tend to show higher contextual fear than males during the first test (p=0.0524, **Fig. 7E**).

These results show that although no significant changes in cued or non-cued fear recall or extinction are observed between sexes following unpaired fear conditioning protocol, females show higher shock reactivity during fear-conditioning, and demonstrate a significantly higher ASR overall during the FPS recall test, regardless of trial type. Notably, this high ASR reactivity observed in females is long-lasting, such as it persists during the second and third FPS test. Lastly, females show significantly higher contextual fear expression than males following the unpaired conditioning protocol.

### 5. Female and male rats show similar FPS following contextual fear conditioning

#### 5.1. FPS after contextual fear conditioning in females and males

In order to compare contextual fear between males and females using FPS without a prior cued fear learning, a separate group of female and male rats were conditioned to context A alone. For direct comparisons, ASR was corrected per 100 g of body weight. Again, females showed a significantly higher shock reactivity than males when corrected per body weight (p=0.0211, **Fig. 8A**). Two-way RM ANOVA showed a significant trial effect of pre-shock vs. post-shock during the first contextual fear recall (F(1, 18)=18.03, p=0.0005), but no significant sex effect (F(1, 18)=0.01802, p=0.6762), nor interaction between sex and trial type (F(1, 18)=0.1052, p=0.7494) were found (**Fig. 8B**). *Post-hoc* analysis showed that ASR during post-shock is significantly higher in comparison to pre-shock in males (p=0.0092), and females (p=0.0249), indicating that both sexes express comparable levels of contextual fear.

**Figure 8.**
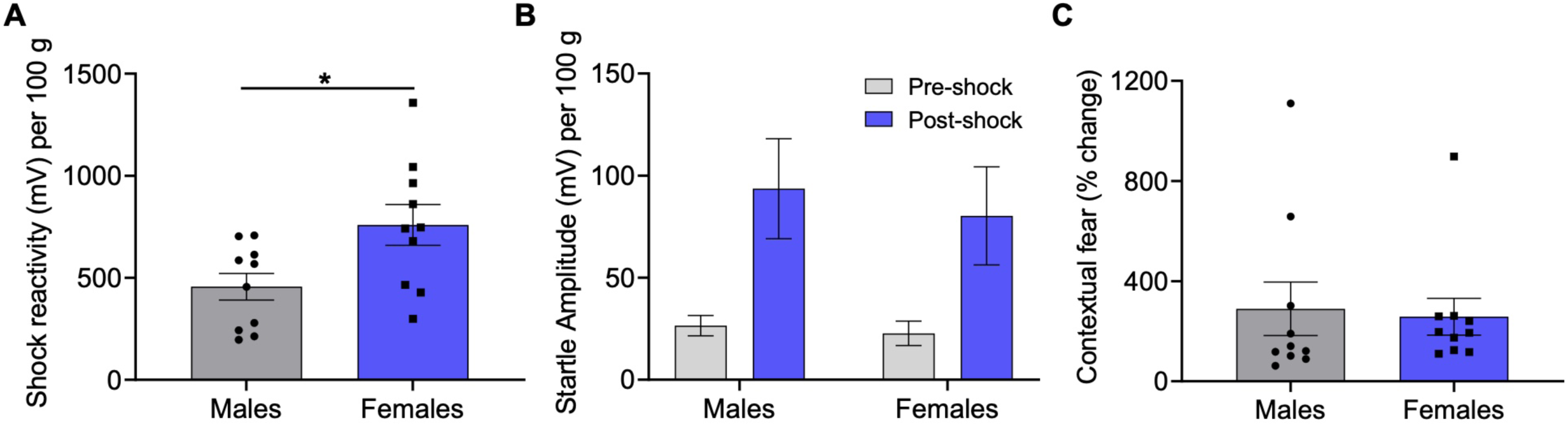
Female and male rats show similar FPS after contextual fear conditioning. **A)** Females show significantly higher shock reactivity than males during contextual fear conditioning. **B)** During the first contextual fear recall, there was a significant trial effect of pre-shock vs. post-shock (p=0.0005), but no significant sex effect (p=0.6762), nor interaction (p=0.7494). **C)** No significant sex differences were found in the contextual fear recall. *p<0.05.

#### 5.2. Percentage change of fear and fear extinction after contextual fear-conditioning in females and males

No significant differences were found in the percentage change of contextual fear between females and males (p=0.8100, **Fig. 8C**). Two-way RM ANOVA showed a significant effect of time on contextual fear extinction (F(1.530, 27.53)=4.722, p=0.0248), but no effect of sex (F(1, 18)=0.3492, p=0.5619), nor interaction between sex and time (F(2, 36)=0.5009, p=0.6101), not shown. These results suggest that males and females express and extinguish contextual fear in a similar fashion.

## Discussion

Our results demonstrate that female and male rats are more alike than different in the classic FPS, showing similar levels of cued, non-cued, and contextual fear expression, despite females having significantly higher shock reactivity during fear-conditioning. However, females show reduced fear discrimination during classic FPS, indicating a parallel reduction of cued fear and increase in non-cued fear recall in comparison to males. Exposure to reduced CS-US contingency, lack thereof, or reversed CS-US order, unmasks more sex differences, with the most striking one following exposure to unpredictable threats (cue and shock unpaired), with females, but not males, showing a long-lasting increase in ASR overall, regardless of trial type, during repeated FPS testing and higher contextual fear expression.

Similar to our results, FPS studies showed no sex differences in the ASR baseline in rats (Russo and Parsons, 2021), but others reported that female rats demonstrate higher ASR than males even when corrected per body weight (Beck et al., 2002; Shanazz et al., 2022). These sex differences can be attributed to decibel level of the ASR, the estrus cycle, and the previous exposure to a stressor (Beck et al., 2008). In contrast to our results, female rats demonstrate reduced reactivity to foot-shocks than males (Russo and Parsons, 2021). However, shock reactivity in the study was measured by computing motion - averaging pixel changes per frame, whereas in the current study we measured the amplitude of the whole-body jump during each foot-shock. In agreement with our findings, studies reported higher shock reactivity in female rats during fear-conditioning (Mitchell et al., 2022), and higher sensitivity to foot-shocks shown as females’ lower thresholds to jump (Beatty and Fessler, 1977; for review see: van Haaren et al., 1990). These sex differences in the reactivity to foot-shock might be also due to differences in the stress axis and the sympathetic nervous system reactivity (Weinstock et al., 1998), or potential differences in pain threshold. For example, human and rodent studies reported that females tend to be more sensitive to nociceptive stimuli (for reviews in humans and rodents see: Mogil and Bailey, 2010; Sorge and Totsch, 2017, respectively).

Prior reports on sex differences in cued fear expression delivered mixed results (for reviews see: Day and Stevenson, 2020; Bauer, 2023). FPS studies in rats (all following fear-conditioning with 10 pairings of foot-shocks and light, as in the current study) reported either no sex differences in cued fear (Zhao et al., 2018), similarly to what we demonstrate in the current study, reduced cued fear (Voulo and Parsons, 2017), or greater cued fear in female in comparison to males rats (de Jongh et al., 2005). Mixed results were also reported when freezing behavior was used as a proxy of cued fear following conditioning to a tone paired with foot-shock, with either no sex differences in rats (Baran et al., 2009; Fenton et al., 2014; Voulo and Parsons, 2017; Urien and Bauer, 2022), reduced cued fear in female rats (Pryce et al., 1999; Baran et al., 2010; Russo and Parsons, 2021), or increased cued fear in female mice (Chen et al., 2014), and rats (Fenton et al., 2016). Notably, although we report no vast sex differences in cued or non-cued fear in the FPS, we demonstrate that females show a reduced fear discrimination index compared to males, which indicates a concomitant reduction of cued fear and an increase in non-cued fear observed in the same female rats. These results are in line with previous preclinical reports showing that female rodents demonstrate reduced fear discrimination between a discrete tone paired with foot-shock (CS+) vs. another tone never paired with foot-shock (safety signal, CS-), in comparison to male rats (Day et al., 2016), and mice (Keiser et al., 2017; Clark et al., 2019), but see (Foilb et al., 2018). Therefore, in addition to deficits in threat vs. safety discrimination, our results suggest that females show higher cued fear generalization than males. Similar results were found in healthy humans, with non-PTSD women showing less fear discrimination between a safe (CS-) vs. threat stimuli (CS+) in comparison to men during both fear acquisition and recall (Lonsdorf et al., 2015). An overgeneralization of fear response to safety signals is an important hallmark of PTSD (for review see: Jovanovic et al., 2012; Kaczkurkin et al., 2017). For example, fear generalization measured with FPS was shown in women diagnosed with PTSD who suffered childhood abuse in comparison with healthy women as controls (Lis et al., 2020). Lastly, these effects might be sex-specific, as studies in children from a highly traumatized urban area reported that girls discriminate less between CS+ vs. CS-in comparison to age-matched boys (Gamwell et al., 2015).

Another hallmark of PTSD is the increased fear reactivity to unpredictable vs. predictable threats. For example, PTSD patients show increased ASR in comparison with generalized anxiety disorder (GAD) patients and healthy controls during the presentation of visual stimuli that were or were not paired with different aversive sounds during FPS, in both women and men combined (Grillon et al., 2009). Notably, PTSD patients did not differ from controls in their responses to predictable threats, such as visual stimuli always paired with aversive outcome (cued fear) (Grillon et al., 2009). Therefore, we next used modified fear-conditioning protocols to mimic unpredictable threat exposure by reducing or removing the CS-US contingency, or changing the CS-US order. Particularly, the fear-conditioning to unpredictable threats (shock and light unpaired), which mimics the human anxiety-potentiated startle protocol (for review see: Grillon and Ernst, 2020), revealed striking sex differences. Here, females show significantly higher ASR overall than males, independently of trial type during the FPS recall (in context B), and this heightened startle persisted through three consecutive FPS tests. This means that following exposure to unpredictable threats, females show an overall heightened and long-lasting state of vigilance, despite the lack of sex differences in ASR at baseline. These findings are in agreement with a recent report from Urien and Bauer (2022) showing higher freezing behavior in context B in females in comparison to males following unpaired fear conditioning. Notably, this increased ASR in the FPS is associated with reduced cued fear in both males and females, which is in agreement with previous studies showing reduced freezing following unpaired vs. paired conditioning in male rats (Goode et al., 2019), male mice (Glover et al., 2020), and both male and female rats (Urien and Bauer, 2022), but see (Baran et al., 2009).

In contrast to cued fear to a discrete stimulus, contextual fear represents fear learning to multiple sensory cues of the environment associated with a threat (Rudy et al., 2004). Interestingly, when conditioned to a context alone (without prior CS-US exposure), we show that males and females express and extinguish contextual fear similarly. Similar levels of contextual fear between male and female rats were shown before with the FPS (Russo and Parsons, 2021), and with freezing behavior as a proxy for fear (Urien et al., 2021). In contrast, higher levels of freezing behavior in threatening contexts were found in male in comparison to female mice (Mizuno et al., 2012) and rats (Poulos et al., 2015; Russo and Parsons, 2021); whereas other studies showed increased contextual fear in female mice (Moore et al., 2010; Keiser et al., 2017). Some of the conflicting findings regarding the level of contextual fear between male and female rats might be related to differences in the experimental design, and more specifically, the timing of the recall following fear-conditioning. Whereas in our study there are no sex differences in rats conditioned to the context alone, we observe differences in the contextual fear recall and extinction (in context A) in rats subjected to prior cued fear conditioning and/or recall. For example, we found that following 100% CS-US contingency protocol, females, but not males, demonstrate higher expression of contextual fear during the second, as opposed to the first, contextual fear testing in context A, which might suggest differences in the duration of fear consolidation and/or generalization. Female mice also showed more contextual fear generalization than males after receiving a single foot-shock in the conditioning context A, and then showed enhanced freezing when exposed to the different context B (Keiser et al., 2017). Lastly, we show that females demonstrate significantly higher contextual fear expression than males when exposed to unpredictable threats (cue and shock unpaired). Overall, these results suggest that females and males show differences in processing unpredictable threats, resulting in higher levels of ASR reactivity and higher levels of contextual fear in females. These findings might aid in our understanding of the neurobiology of unpredictable threat processing in both sexes which might shine a new light on the higher incidence of PTSD in women.

Although cellular mechanisms of sex differences in unpredictable threat processing are outside the scope of the current study, differences in the extended amygdala circuitry might be contributing to these effects. The bed nucleus of the stria terminalis (BNST), part of the extended amygdala, is necessary for the expression of contextual fear (Sullivan et al., 2004; Duvarci et al., 2009; Zimmerman and Maren, 2011), drives fear expression following conditioning to unpredictable threats in both rodents (Goode et al., 2019, 2020), and humans (Clauss et al., 2019; Figel et al., 2019; Naaz et al., 2019), and has been involved in the pathophysiology of PTSD (for review see Miles and Maren, 2019). Recent reports also suggest potential sex differences in the BNST contributions to contextual fear and fear responses to unpredictable threats. Although excitotoxic lesions of the entire BNST in rats disrupted contextual fear in both sexes (Urien et al., 2021), contextual fear expression led to higher cFos expression (Urien and Bauer, 2022), and upregulation of activity-regulated cytoskeleton-associated protein (Arc) in the anterolateral BNST of male but not female rats (Urien et al., 2021). These results suggest sex differences in the BNST activation during contextual fear expression. Notably, increased cFos expression in the BNST was also found during fear recall in both sexes following unpaired fear-conditioning protocol, despite females showing higher level of fear following this protocol (Urien and Bauer, 2022). The BNST involvement in the processing of unpredictable threats was also demonstrated with a backward conditioning protocol when a neutral cue occurred immediately after the offset of the foot-shock (Goode et al., 2019). This study reported reduced freezing behavior during cued fear recall following backward conditioning in male rats, which we confirm in the current study using FPS. However, we also show that females only tend toward a reduction in cued fear, whereas both sexes show significantly higher contextual fear expression following backward vs. forward cued fear conditioning. Higher levels of contextual fear following reduced or absent CS-US contingency are in agreement with a competitive model of the associative learning (Marlin, 1981).

In addition to variances in the underlying neurocircuitry, some of the reported sex differences in fear expression might result from differences in the behavioral strategies, such a startle vs. freezing, as clearly shown by Russo and Parsons (2021), and other sex-specific fear behaviors such as darting (Mitchell et al., 2022). Freezing is the most commonly used proxy of learned fear in rodents (Maren, 2001; Rudy et al., 2004; Sun et al., 2020), whereas ASR and FPS are used in both animal (de Jongh et al., 2005; Voulo and Parsons, 2017; Zhao et al., 2018; Martinon et al., 2019), and human studies (Grillon et al., 2009; Lis et al., 2020). During the FPS, the rapid nature of the ASR reflex (less than 200 msec from the ASR-eliciting noise onset), allows a precise temporal resolution to dissect out both cued and non-cued components of fear recall. In contrast, the persistent nature of freezing behavior, which often surpasses presentation of a discrete cue, might hamper the ability to distinguish subtle differences in fear measures during and between cue presentations. Furthermore, in contrast to the lack of freezing without prior conditioning or without the presence of CS+/or context, FPS measures the magnitude of fear recall and extinction based on ASR baseline from individual rats and humans. Non-cued fear, first reported by Walker and Davis (2002), reflects potentiation of the ASR during FPS recall, which begins and then persists beyond the first and the following cue presentations (Walker and Davis, 2002). Previous studies from the Rosen group suggested that this FPS component represents a form of anticipatory fear or ‘background anxiety’ (Missig et al., 2010; Ayers et al., 2011). To avoid a conceptual confusion regarding typically measured anxiety responses, in our current and previous studies (Moaddab and Dabrowska, 2017; Martinon et al., 2019), we refer to this FPS component as a non-cued fear. The Rosen group demonstrated that background anxiety/non-cued fear is independent from contextual fear, because it is specifically activated by the cue (CS+) in a modified and not original context (Missig et al., 2010; Ayers et al., 2011). Furthermore, non-cued fear does not appear immediately during fear recall, in so called post-shock ASR (10 initial ASR trials before cue presentation), which conceivably argues against a major contextual fear contamination (Missig et al., 2010, Moaddab and Dabrowska, 2017). Therefore, to fully separate non-cued fear from contextual fear, we postulate that the non-cued fear should be calculated using the post-shock, and not pre-shock ASR, as a baseline (**Fig. 1F**). Lastly, we demonstrate that all modified fear-conditioning protocols (backward, reduced contingency, CS-US unpaired), which lead to reduced cued fear formation, also reduce the non-cued fear in context B, whereas they significantly increased contextual fear, when tested in context A on the next day. Notably, in both females and males, non-cued fear does not extinguish at the same pace as cued fear. These results further demonstrate that non-cued fear depends on efficient cued fear formation and represents cued fear generalization, which is mostly independent of contextual fear. Yet, different cellular mechanisms might be responsible for shifting a balance between cued vs. non-cued fear formation (Moaddab and Dabrowska, 2017; Janeček and Dabrowska, 2019; Martinon et al., 2019).

This study has some limitations, such as the use of naturally cycling females without monitoring their estrous cycle. Although previous reports showed that female hormones do not affect the expression or extinction of the FPS (Zhao et al., 2018; Voulo and Parsons, 2019), other reports showed that females fear-conditioned during proestrus show reduced FPS during the second half of fear recall test similarly to males, but females that were fear conditioned during late diestrus showed higher ASR vs. males and vs. females conditioned during proestrus, yet these findings were not replicated when measuring freezing behavior in the same study (Carvalho et al., 2021) but see (Gruene et al., 2015). A recently published meta-analysis of fear responses in female and male rodents shows that estrous cycle does not contribute to increased variability in females datasets in comparison to males (Kaluve et al., 2022). Nonetheless, as PTSD pathophysiology in human females have been linked to reproductive hormones (Ravi et al., 2019), future studies should focus on investigating whether a similar link exists in rodent females in response to unpredictable threats.

Preclinical studies studying the neurobiology of fear processing in female rodents are scarce (Shansky and Murphy, 2021; Bauer, 2023), and most of these studies used freezing behavior as a proxy of fear. Here, we used FPS, which offers precise temporal resolution needed to dissect out different fear outcomes over time, and allows translational comparisons between rodents and humans. After exposure to unpredictable threats, females showed a heightened and long-lasting state of vigilance and higher levels of contextual fear than males. These sex differences in unpredictable threat processing might help to better understand the neural mechanisms underlying the higher incidence of PTSD in women. The modified FPS protocols used in this study may be incorporated into future preclinical studies focusing on potential pharmacological targets to treat PTSD in men and women.

## Acknowledgements

We are very thankful to Rachel Chudoba, Susan H. Olson, Andrew Forrest, Rachel Parker, and Amaan Azeemullah for their help with behavioral tests. We also thank Rachel Chudoba for proofreading the manuscript.

